# Genomic insights into hybrid zone formation: the role of climate, landscape, and demography in the emergence of a novel hybrid lineage between *Populus trichocarpa* and *P. balsamifera*

**DOI:** 10.1101/2023.07.17.549358

**Authors:** Constance E. Bolte, Tommy Phannareth, Michelle Zavala-Paez, Brianna N. Sutara, Muhammed F. Can, Matthew C. Fitzpatrick, Jason A. Holliday, Stephen R. Keller, Jill A. Hamilton

## Abstract

Population demographic changes, alongside landscape, geographic, and climate heterogeneity, can influence the timing, stability, and extent of introgression where species hybridize. Thus, quantifying interactions across diverged lineages, and the relative contributions of interspecific genetic exchange and selection to divergence at the genome-wide level is needed to better understand the drivers of hybrid zone formation and maintenance. We used seven latitudinally arrayed transects to quantify the contributions of climate, geography, and landscape features to broad patterns of genetic structure across the hybrid zone of *Populus trichocarpa* and *P. balsamifera* and evaluated the demographic context of hybridization over time. We found genetic structure differed among the seven transects. While ancestry was structured by climate, landscape features influenced gene flow dynamics. Demographic models indicated a secondary contact event may have influenced contemporary hybrid zone formation with the origin of a putative hybrid lineage that inhabits regions with higher aridity than either of the ancestral groups. Phylogenetic relationships based on chloroplast genomes support the origin of this hybrid lineage inferred from demographic models based on nuclear data. Our results point towards the importance of climate and landscape patterns in structuring the contact zones between *P. trichocarpa* and *P. balsamifera* and emphasize the value whole genome sequencing can have to advancing our understanding of how neutral processes influence divergence across space and time.

## Introduction

Understanding the processes influencing the formation and maintenance of species is a central goal of evolutionary biology and is crucial to the management and conservation of biodiversity in a rapidly changing environment (Frankham, 2010; Hoffmann et al., 2015; Payseur & Rieseberg, 2016). As such, hybridization and introgression have long been of interest (Dobzhansky, 1936; Heiser, 1949; Jeffrey, 1916; Stebbins, 1959), and recent advances in population-scale genome sequencing have revolutionized our understanding of the frequency and pervasiveness of hybridization in nature (Hamilton & Miller, 2016; Janes & Hamilton, 2017; VanWallendael et al., 2022). Multiple taxa are known to form natural hybrid zones; however, they are particularly prevalent among forest tree species (Abbott, 2017; Suarez-Gonzalez et al., 2018; Swenson & Howard, 2004). Climatic oscillations during glacial and interglacial periods have influenced the distributions of many temperate and boreal trees, facilitating opportunities for interspecific gene flow during times of secondary contact (Hamilton et al., 2015; Hewitt, 2004; Jump & Peñuelas, 2005; Soltis et al., 2006; Swenson & Howard, 2004). However, despite widespread observations of hybridization across forest tree species, gaps remain in our understanding of how demographic, neutral and nonneutral evolutionary processes influence the formation and maintenance of natural hybrid zones in forested landscapes across space and time. Teasing apart these processes requires an assessment of the historical, landscape, and climatic factors that underlay hybrid zone formation. Ultimately, understanding how population demographic changes, alongside landscape heterogeneity and climate adaptation contribute to the timing, stability, and extent of hybridization will be critical to managing species in a rapidly changing climate.

Population demographics, including expansion and contraction of a species’ range can have substantial influence on standing genetic variation, impacting the evolutionary trajectory of populations across space and time. Hybrid zones often form when previously isolated lineages come into contact and the impacts of these processes, including genetic drift and gene flow, likely play an important role shaping the genetic structure of a contact zone (Abbott, 2017; N. H. Barton & Hewitt, 1989). Shifts between isolation and contact can coincide with changes in effective population size (*N*_e_) influencing the direction and extent of genetic exchange, while fine-scale variance at the population or genome-level can further influence elimination or fixation of alleles. In forest trees, genetic exchange often appears to be largely asymmetrical favoring the genomic background of one species due solely or in part to differential dispersal capacity, wind-patterns, and unidirectional reproductive incompatibilities (El Mujtar et al., 2017; Hamilton et al., 2013a; Lepais et al., 2009; Lexer et al., 2005, 2006). Given that an influx of genetic variants via interspecific gene flow not only buffers *N*_e_, but can generate novel genetic recombinants upon which natural selection can act, characterizing genetic structure and the impact genetic exchange may have across space and time is needed to predict evolution.

Demographic inference can be particularly useful for characterizing processes underlying patterns of genetic exchange that can influence species evolutionary relationships, distributional shifts, or adaptive potential (Bacilieri et al., 1996; Petit et al., 2004). Several forest tree hybrid zones have characterized patterns of introgression, including oaks (Cannon & Petit, 2020; Eaton et al., 2015; McVay et al., 2017), poplars (Chhatre et al., 2018; Christe et al., 2017; Lexer et al., 2005; Suarez-Gonzalez et al., 2016; Suarez-Gonzalez et al., 2018), and spruce (Hamilton et al., 2013a, 2013b). However, few studies have leveraged whole genome sequencing to tease apart the temporal and spatial dynamics underlying the evolutionary history of contact zones using both biparentally and uniparentally inherited genomes. Viewing distinct evolutionary trajectories associated with biparentally and uniparentally inherited genomes through the lens of demographic change and landscape heterogeneity enables assessment of the role these evolutionary processes may play to the formation and maintenance of natural hybrid zones.

Here, we use *Populus*, an ecologically and economically important model system in forest trees that naturally hybridize with congeners where geographical ranges overlap. *Populus trichocarpa,* the first tree to have its genome fully sequenced (Tuskan et al., 2006), has played an extensive role in our understanding of tree genome biology, comparative genomics, and adaptive introgression (e.g., Jansson & Douglas, 2007; Shang et al., 2020; Suarez-Gonzalez et al., 2016). Natural hybrid zones exist between *P. trichocarpa* and *P. balsamifera*, largely associated with geographically and climatically steep transitions from maritime to continental climates west and east of the Rocky Mountains. However, divergence has likely involved a combination of intrinsic and extrinsic factors, including a dynamic history of population size change throughout glacial and interglacial periods, landscape-related barriers to gene flow, and climatically-related responses to selection (Geraldes et al., 2014; Keller et al., 2010; Levsen et al., 2012; Slavov et al., 2012). We compare whole-genome sequence data for biparentally (nuclear) and uniparentally (chloroplast) inherited genomes to quantify divergence and the evolutionary relationship between species across space and time. This will provide an understanding of the role neutral and non-neutral processes play in the formation of these long-lived hybrid zones (Petit et al. 2005).

Using a sampling design of latitudinally arrayed transects, we sequenced whole genomes for 576 trees to capture parental-types and hybrids across repeated zones of contact between *P. trichocarpa* and *P. balsamifera* distributed across their entire range of overlap. With these data, we observed genetic structure within and across each contact zone and asked (1) How has climate, geography, and landscape-level barriers influenced genetic structure, and (2) What is the history and extent of interspecific gene flow? These data provide new insights into how different evolutionary processes contribute to the formation and persistence of hybrid zones over varying spatial and temporal scales.

## Materials and Methods

### Sampling of plant material

In January 2020, vegetative branch cuttings were collected from 576 poplar trees spanning repeated natural contact zones between *Populus trichocarpa* and *Populus balsamifera* (Figure 1a). These represent individual collections from seven latitudinally distributed transects (hereafter referred to as Alaska_1, Alaska_2, Cassiar, Chilcotin, Jasper, Crowsnest, and Wyoming) spanning much of the species’ overlapping distributions, from 40°N to 65°N and −100°W to −150°W longitude. On average the distance between samples was approximately 43 kilometers (range: <1 - 962 km). Vegetative branch cuttings were transported back to a greenhouse at Virginia Tech (Blacksburg, VA, USA) for propagation. Details associated with propagation and greenhouse conditions are provided in Appendix 1 of Supplementary.S1. Young leaf tissue was sampled from each of the 576 propagated branch cuttings, placed immediately on dry ice, and then transferred to a −80°C freezer for long-term preservation.

**Figure 1.**
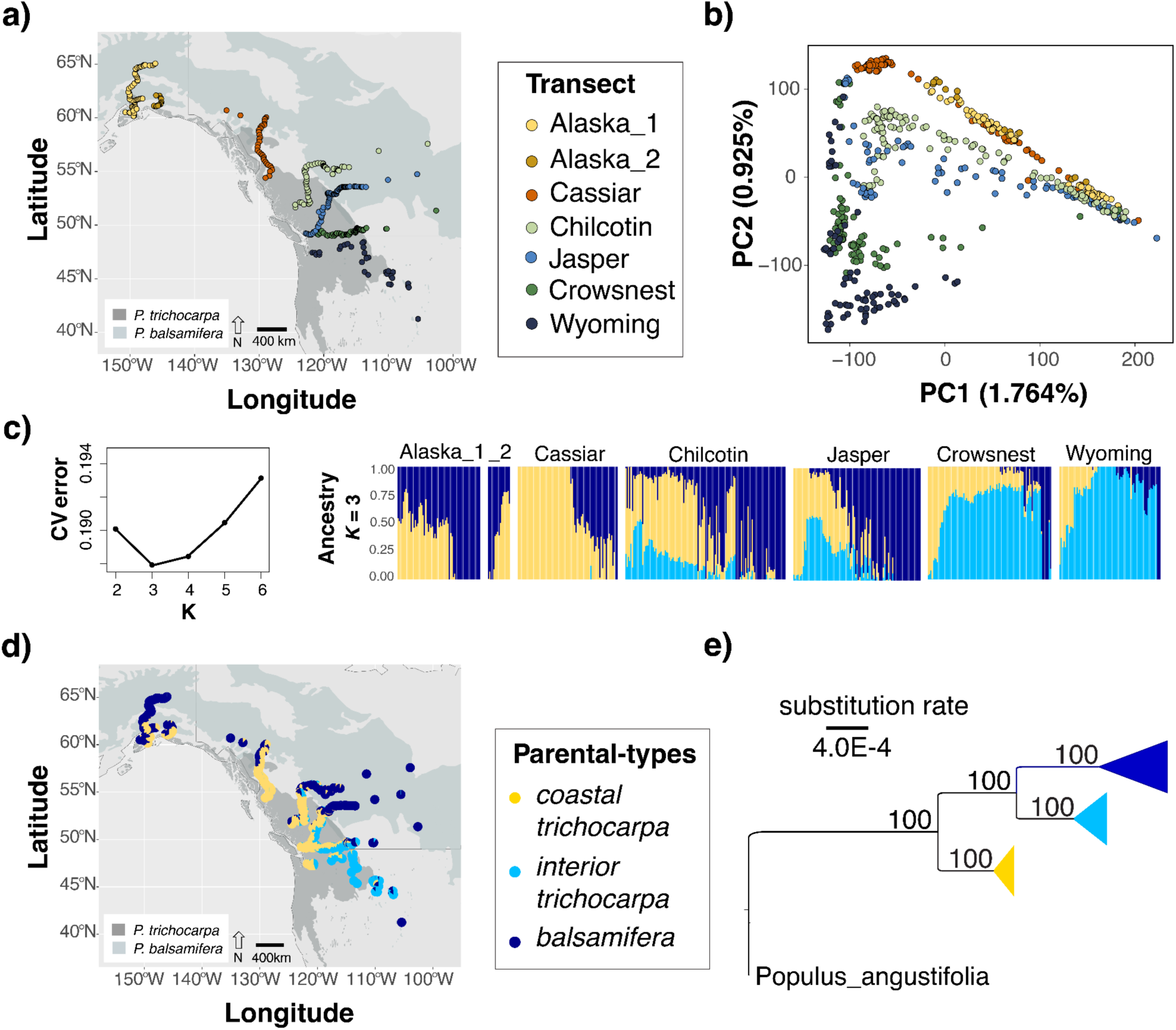
Spatial patterns of genetic structure and ancestry across 546 trees sampled from a) seven transects of the *Populus balsamifera* and *P. trichocarpa* hybrid zone, labeled by transect; b) transect-specific genomic structure based on Principal Component Analysis for PC1 and PC2 c) results of admixture analysis for each sample at *K* = 3, the best supported cluster assignment on the basis on lowest cross validation (CV) error; d) geographic distribution of *K =* 3 ancestry proportions illustrated as pie charts; e) ML phylogeny of chloroplast evolutionary relationships among parental-type samples from *K =* 3 admixture results (Q > 0.90 for each cluster). Bootstrap confidence is provided on each branch.

### Genomic Library Preparation and Bioinformatics

For each of the 576 samples, DNA was extracted from approximately 100mg of leaf tissue using the Qiagen plant DNeasy kit, with one modification (details available in Supplementary.S1; Appendix 2). Genomic DNA libraries were constructed at the Duke University Center for Genomic and Computational Biology, using an Illumina DNA Prep kit (Illumina Inc., San Diego, USA). Genomic libraries were sequenced on an S4 flow cell in 2×150bp format on an IlluminaNovaSeq 6000 instrument with 64 samples per lane. De-indexing, QC, trimming adapter sequences, and sequence preprocessing were completed by the sequencing facility.

Subsequent bioinformatic analysis was performed on Virginia Tech’s Advanced Research Computing System (ARC) using Burrows-Wheeler Aligner (BWA) to map reads to the *P. trichocarpa* reference genome (v4.0), and the resulting SAM files were converted to BAM format with SAMtools (Li & Durbin, 2010). The Genome Analysis Toolkit (v3.7) Haplotype Caller algorithm was then used to generate individual gVCF files, which were merged in a single VCF file with the GATKGenotype GVCFs function. This raw VCF had ∼82 million variant calls and was quality-filtered for variants that had poor map quality (MQ < 40.00), elevated strand bias (FS > 40.000, SOR > 3.0), differential map quality between reference and alternative alleles (MQRankSum < - 12.500), positional bias between reference and alternate alleles (ReadPosRankSum < - 8.000), or low coverage depth (QD < 2.0).

After quality filtering, ∼34 million single nucleotide polymorphisms (SNPs) and ∼4.8 million indels remained in the variant call set. Filtering removed all of the ∼28 million multiallelic sites (defined as having more than two allelic states), of which ∼5 million were SNP sites (i.e., did not include indels). For 575 of the 576 samples, missing data was on average 0.7 % (SD: 0.1 - 1.3 %). One sample, GPR-14_S50_L001, had 99.9% missing data and was therefore not retained for further analysis.

### Analysis of population structure

To account for potential sampling of non-focal poplar species and remove those samples, we merged the genetic data for our 575 samples with genetic data available for 48 samples of *P. angustifolia*, which overlaps in portions of the southern distribution of our sampling region (Chhatre et al., 2018). Methods in Supplementary.S1 (Appendix 3) detail the analyses used. While we found no need to remove samples due to admixture with *P. angustifolia*, we removed 29 of the 575 samples with unknown ancestry (Supplementary.S1; Figure S1 and S2), leaving 546 samples for subsequent genetic analyses.

To assess the genetic structure of the contact zones between *P. balsamifera* and *P. trichocarpa*, we first thinned variant sites in high linkage disequilibrium using *Plink* (v1.9) based on 10,000 bp windows sliding in increments of 1000 bp along each chromosome, flagging site pairs with an *r*^2^ > 0.1. One site for each pair exceeding the *r*^2^ threshold (i.e., ‘prune.out’ file), along with singletons (--mac 2) and those with missing data (--max-missing 1.0), were removed using *vcftools*, version 0.1.16. The resulting 299,335 low LD variant sites were used to visualize the scale and spatial extent of hybridization. Genomic structure was assessed using a principal component analysis (PCA) with the ‘prcomp’ function of the *stats* package in R, version 4.2.1. We then performed an ADMIXTURE analysis (Alexander et al., 2009) using cluster assignments ranging from *K*=2 to *K*=6. The cluster assignment (*K*) with the lowest cross validation error (k-folds = 10) was determined to best fit the data (Figure 1c).

### Assessing evolutionary relationships from nuclear data

To evaluate the evolutionary relationships between genetic clusters, we used the three genetic clusters identified from ADMIXTURE (*K* = 3; see Results) to determine the topology with the greatest support. We identified parental-type samples for each genetic cluster, hereafter referred to as *P. balsamifera*, *coastal P. trichocarpa*, and *interior P. trichocarpa*, based on K=3 using an ancestry coefficient threshold of Q > 0.90. Of the 546 samples included in the analysis, 185 parental-types were identified (112 *P. balsamifera*, 51 *coastal P. trichocarpa*, and 22 *interior P. trichocarpa*). Using parental-types, we evaluated three different topologies based on the proposed divergence history between *interior P. trichocarpa* and the other two parental-type genetic lineages (Table 1). The first topology evaluated expectations based on a recent divergence from *coastal P. trichocarp*a, while the second tested for a recent divergence from *P. balsamifera*, and the third evaluated whether *interior P. trichocarpa* was a hybrid lineage formed from an admixture event between *coastal P. trichocarpa* and *P. balsamifera*.

**Table 1.** Demographic inference AIC comparisons across 3 population models. Model abbreviations are defined in the footnote. The lower the AIC and delta AIC the better the model fit to the data. The first three rows (Table 1a) are results from the topology test to identify the evolutionary relationship of *interior P. trichocarpa* (I) to *P. balsamifera* (B) and *coastal P. trichocarpa* (C). The following four rows (Table 1b) are results of four models with gene flow parameters tested in various scenarios under the best fit topology (Topology 3; Figure S6) of admixed origin. Bolded models in each of the two inference routines signify best fit with delta AIC = 0.00.

| **a)**<br>**Model** | **k** | **AIC** | **delta**<br>**AIC** |
| --- | --- | --- | --- |
| Topology 1: SI | 6 | 68,945.17 | 4,942.96 |
| Topology 2: SI | 6 | 75,772.18 | 11,769.97 |
| Topology 3: SI | 6 | 64,002.21 | **0.00** |

| **b)**<br>**Model** | **k** | **AIC** | **delta**<br>**AIC** |
| --- | --- | --- | --- |
| Topology 3: T2_SC | 12 | 34,935.00 | 136.26 |
| Topology 3: T2_IM | 12 | 43,391.55 | 8,592.80 |
| Topology 3: T3_IM_SC | 14 | 34,798.75 | **0.00** |
| Topology 3: T4_SC_SC | 14 | 35,426.09 | 627.34 |

\*Models; SI (strict isolation), IM (isolation-with-migration), SC (secondary contact), T# (number of time intervals considered)

The evolutionary relationship of the three genetic clusters was evaluated using the genotypes from the 185 parental-types (112 *P. balsamifera*, 51 *coastal P. trichocarpa*, and 22 *interior P. trichocarpa*) and Diffusion Approximation for Demographic Inference (*∂α∂i* v.2.1.1;(Gutenkunst et al., 2009). We used the unfolded joint site frequency spectrum (jSFS) of derived allele counts following polarization of genotypes following the approach of (Luqman et al., 2023). Details pertaining to data sources and program implementation are provided in Supplementary_S1, Appendix 4. We assigned allelic states as either ancestral or derived based on the consensus ancestral state: 6495 sites (2.260%) had the reference allele assigned to the derived state and 275,751 sites (95.959%) had the reference allele assigned to the ancestral state. The remaining 1.781% of sites had undefined ancestral alleles. To increase computational efficiency, we used the derived allele frequencies for a random sample of 10 diploid individuals per parental-type cluster to build an unfolded 3-dimensional joint site frequency spectrum (jSFS) using the function ‘GenerateFs’ in the dadi-cli program (X. Huang et al., 2023). We performed ten replicate runs of each model in *∂α∂i* with a 40 x 50 x 60 grid space and the nonlinear Broyden-Fletcher-Goldfarb-Shannon (BFGS) optimization routine. Akaike information Criterion (AIC; Akaike 1974) was employed for model selection, with the best replicate run (highest log composite likelihood) for each model used to calculate ΔAIC (AIC_model i_ – AIC_best model_) scores (Burnham & Anderson, 2002).

To estimate the timing of divergence and quantify the extent and directionality of gene flow between genetic clusters, we tested four hypotheses, or models, of connectivity and divergence based on the best supported topology (Table 1). The first model assumed a scenario of recurrent gene flow, while the other three models assumed scenarios of isolation that varied based on the timing and frequency of secondary contact (Supplementary.S1, Figure S4). We performed ten replicate runs of each model in *∂α∂i* with a 40 x 50 x 60 grid space and the nonlinear Broyden-Fletcher-Goldfarb-Shannon (BFGS) optimization routine. Akaike information Criterion (AIC; Akaike 1974) was employed for model selection, and Fisher Information Matrix (FIM)-based uncertainty analysis was then conducted for the best supported model to obtain upper and lower 95% confidence intervals (CIs) for the inferred parameter estimates. To unscale parameter estimates and their 95% CIs we used a mutation rate of 1.33 x 10^-10^ substitutions/site/year rate for *Populus trichocarpa* (Hofmeister et al., 2020), a generation time of 15 years, and a calculated effective genome length (Supplementary.S3). To calculate effective genome length, we multiplied the proportion of variant sites used in the modeling (count of filtered sites in each pairwise inference with no missing data and singletons removed / count of variant sites prior to filtering) by the number of base pairs from the *P. trichocarpa* reference genome considered in variant calling (389,204,664 bp).

### Assessing evolutionary relationships from chloroplast data

To gain insight into the evolutionary relationships among the three genetic clusters identified from nuclear data, we used the 185 parental-types to examine differentiation across uni-parentally inherited chloroplast genomes. Chloroplast (cp) genomes were assembled using NOVOPlasty (Dierckxsens et al., 2017). The published Nisqaully-1 *P. trichocarpa* chloroplast genome (NC_009143.1) was used as the reference sequence for genome assembly. The ribulose-bisphosphate carboxylase (*rbc*L) gene was manually extracted and used as a seed, serving as an initial reference point for the chloroplast genome assembly. Assembled chloroplast genomes were annotated using Geneious v7.1.4 (Dierckxsens et al. 2016). Following annotation, inverted repeats and coding regions were manually checked for each annotated individual. In total, 184 out of the 185 parental-types were successfully assembled, with one individual (genotype 831) excluded due to incomplete assembly.

A maximum likelihood (ML) phylogenetic tree was built to assess the relationships among the chloroplast genomes for each parental type. The chloroplast genomes from two *P. angustifolia* samples (NC_037413 and MW376761) were downloaded from NCBI and selected as an outgroup. Chloroplast genomes for all 184 samples and outgroups were aligned using MAFFT V.7.453 (Katoh & Standley, 2013) to perform a ML phylogeny in IQ-tree 1.6.11 (Trifinopoulos et al., 2016). The best fit evolutionary model for the sequence alignment was calculated by ModelFinder (Kalyaanamoorthy et al., 2017), with the selected model being the transversion model with empirical base frequencies and FreeRate model (TVM+F+R2). A ML tree was generated following 1,000 bootstrap replications and a consensus phylogenetic tree was created based on all parental-type samples using the FigTree software v1.4.13.

### Associating genetic structure with climate and geography

To quantify the multivariate relationships between genetic structure, climate, and geographic variation across the hybrid zone, a redundancy analysis (RDA) was conducted within the *vegan* package, version 2.6-2 (Oksanen et al., 2020) in R version 4.2.1. Climate data at 30 second resolution averaged across the years 1961-1990 for twenty-five variables associated with geographic origin were extracted using *ClimateNA* (version 5.2; (Wang et al., 2016) for each sampled tree (Supplementary.S2) and then tested for collinearity using variance inflation factor (VIF) analysis implemented in the *usdm* R package (version 1.1-18; (Naimi et al., 2014). Six climatic variables with a VIF less than 10 were retained for subsequent analysis, including continentality (TD), mean annual temperature (MAT), mean annual precipitation (MAP), climate moisture deficit (CMD), relative humidity (RH), and precipitation as snow (PAS). Geographic variables included for subsequent analysis were latitude, longitude, and elevation.

Genotype data (299,335 nuclear SNPs and 2,828 chloroplast SNPs) for all 546 samples was converted to minor allele counts per sample using vcftools (version 0.1.16). A permutation-based analysis of variance (ANOVA) procedure with 999 permutations (Legendre & Legendre, 2012) was used to test the overall statistical significance of the RDA model (*α* = 0.05) as well as the significance of each RDA axis separately. Variance partitioning was then performed using the ‘varpart’ function of the *vegan* package to quantify the influence of predictor variables and their confounding effects to observed genetic variation. This analysis was repeated without hybrid samples (0.10 < Q < 0.90) to compare the multivariate relationships among the 185 parental-type samples.

### Visualizing barriers to gene flow

To evaluate genetic connectivity across the contact zones between *P. trichocarpa* and *P. balsamifera,* we used EEMS (Estimated Effective Migration Surfaces; (Petkova et al., 2016) and observed correspondence between landscape features, ancestry, and patterns of genetic connectivity. A genetic dissimilarity matrix was generated using the ‘bed2diffs script’ included within the EEMS program informed by the 299,335 genome-wide nuclear SNPs across 546 individuals. Two additional geographic files were created using geographic coordinates; including the location data of each individual (.coord extension) and a large general habitat polygon (.outer extension) which was manually obtained using Google Maps API v3 Tool. The geographical coordinates for each sample alongside the parameterized grid space (500 demes with each grid cell ∼21,000 km^2^) assigned individuals to one of 49 demes, defined as pseudo populations of close geographical proximity to a vertex of the defined grid space (Supplementary.S1, Figure S3). We then estimated the effective migration rate using a stepping-stone model. Model parameters included a burn-in of 25,000,000 with an MCMC length of 50,000,000 and thinning to sample 1 in every 100,000 iterations. The results of three replicate runs were averaged to visualize effective diversity (q) and effective migration rate (m) surfaces (in log_10_ scale) across the contact zone using the R package *rEEMSplots* v 0.0.1 (Petkova et al., 2016).

## Results

### Genomic structure of hybrid zone identifies three distinct genetic clusters

Using all 546 individuals, the nuclear genomic PCA ascribed genetic structure along PC1 (1.764% variance explained) to species-specific differences between *P. trichocarpa* (far left of PC1) and *P. balsamifera* (far right of PC1) and PC2 (0.925% variance explained) to genetic structure within *P. trichocarpa* (Figure 1b). Admixture analysis best supported the presence of three genetic clusters (*K =* 3), with admixed individuals present across all transects (Figure 1c). Two of the three genetic groups clustered geographically within the range of *P. trichocarpa* (Little 1971) – this included a *coastal P. trichocarpa* cluster which had a northern, coastal distribution, and an *interior P. trichocarpa* cluster, which had a southern, interior distribution (Figure 1d). The third genetic cluster was *P. balsamifera* in the K = 3 admixture analysis. Pairwise estimates of genome-wide genetic differentiation (*F*_ST_) were similar across the three genetic clusters, ranging from 0.053 for *coastal - interior P. trichocarpa,* 0.054 for *interior P. trichocarpa - P. balsamifera*, and 0.056 for *coastal P. trichocarpa - P. balsamifera* comparisons.

Of the 546 samples included in the analysis, 185 parental-types associated with *K* = 3 genetic clusters were identified based on Q > 0.90 (112 individuals identified as *P. balsamifera*, 51 individuals identified as *coastal P. trichocarpa*, and 22 individuals identified as *interior P. trichocarpa). Coastal P. trichocarpa* and *P. balsamifera* genetic ancestry was observed in Alaska and Cassiar transects, although the Alaska transect did not appear to have any *coastal P. trichocarpa* parental-types. All three genetic clusters were observed in Chilcotin, Jasper, Crowsnest, and Wyoming transects (Figure 1c). Shared ancestry associated with *P. balsamifera* along the Crowsnest transect was restricted to the eastern edge of the *P. trichocarpa* distribution. The Wyoming transect was dominated by *interior P. trichocarpa* parental-type ancestry (Q > 0.90 at *K* = 3; Supplementary.S2), whereas individuals sampled near the western edge of this transect exhibited mixed ancestry between the coastal and interior genetic clusters of *P. trichocarpa*. Consistent with the Crowsnest transect, *P. balsamifera* ancestry was limited to the far eastern distribution of *P. trichocarpa,* with *P. balsamifera* parental-types restricted to fragmented populations geographically distant from the main distribution of either species.

### Nuclear and chloroplast genomes identify the persistence of three genetic lineages

Given the similar estimates of genome-wide nuclear differentiation (*F*_ST_), we performed a topology test to better understand the relationships among the three genetic clusters. The best fit topology as determined by AIC (Table 1a) supported *interior P. trichocarpa* as an ancient lineage of admixed origin. The second-best fit topology placed *interior P. trichocarpa* as a split from *coastal P. trichocarpa*, but delta AIC was 4,943 units higher.

To complement assessment of nuclear topologies, evolutionary relationships were also assessed using ML analysis of chloroplast genomes. The alignment of the 184 individuals and outgroup genomes produced a single matrix with 164,183 nucleotide sites, including 1,020 polymorphic sites. Three distinct clades were detected using the ML tree, including *coastal P. trichocarpa*, *interior P. trichocarpa*, and *P. balsamifera* respectively. The *coastal P. trichocarpa* clade appeared ancestral to both *interior P. trichocarpa* and *P. balsamifera*, with all three exhibiting robust bootstrap support. Lineage assignments were congruent between nuclear and chloroplast genomes, except for individual 757, which clustered with *interior P. trichocarpa* in the nuclear admixture analysis but in the chloroplast ML tree belonged to the *P. balsamifera* clade.

### Climate strongly associates with genetic structure

Climate and geography explained 3.79% (*r*^2^) of the genetic variance across the 546 sampled trees and 299,335 SNP dataset. Most of the explainable genetic variance (PVE) was accounted for by the first RDA axis (38.26%, Figure 2a) with continentality (TD) and mean annual temperature (MAT) having the highest predictor loadings (eigenvectors). Individuals identified as *P. balsamifera* were associated with higher TD and lower MAT on average. The second RDA axis captured 19.23% (PVE) of the explainable genetic variance, with the highest predictor loadings on the second RDA axis being relative humidity (RH), elevation, and longitude. These variables were associated with differentiation within *P. trichocarpa* across the seven transects. Associations between climate, geography and genetic variation were not observed along RDA2 for *P. balsamifera* (Figure 2). Combining loadings for RDA1 and RDA2, we found latitude, TD, and climate moisture deficit (CMD) had the strongest associations with genetic variation within and among sampled transects.

**Figure 2.**
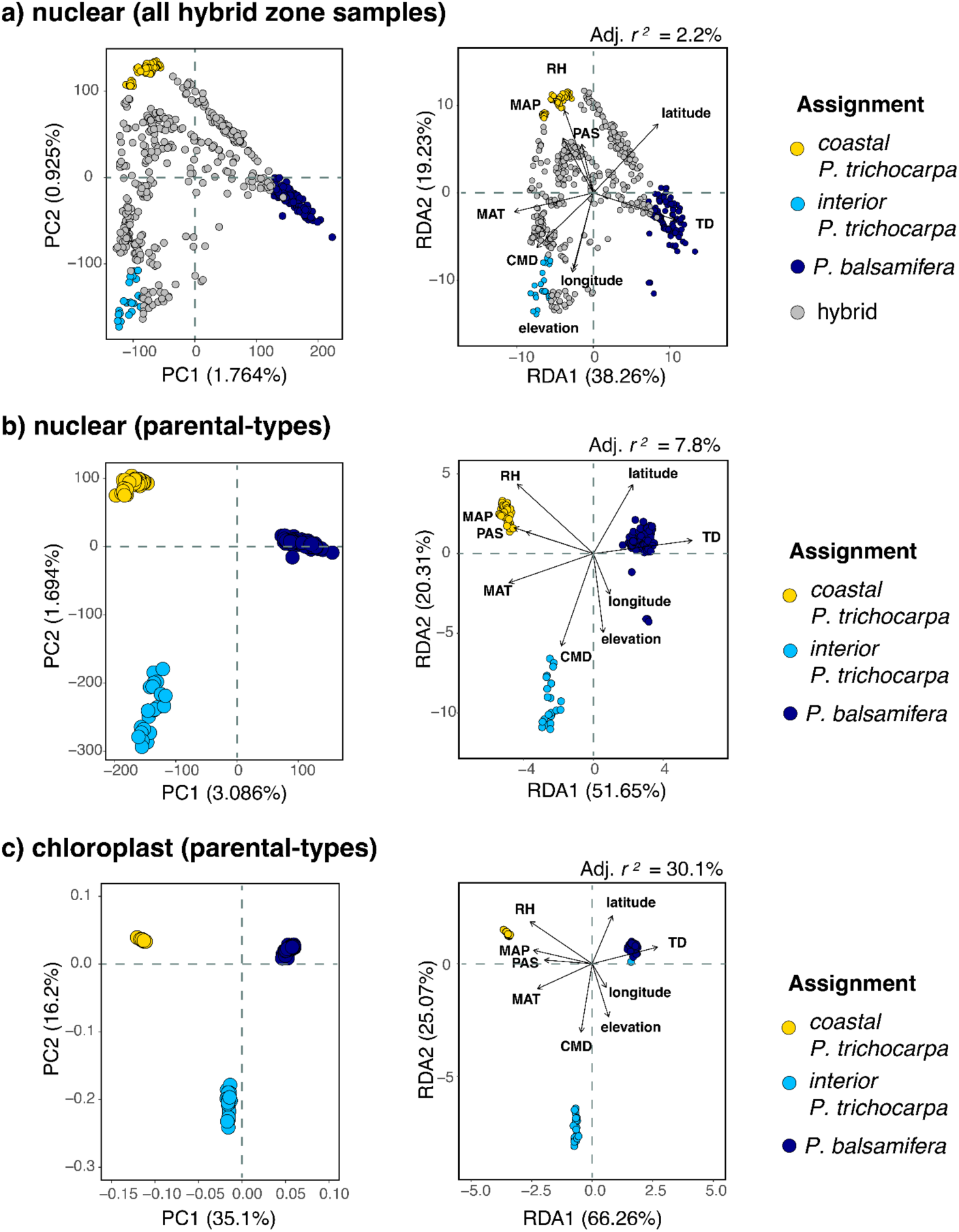
Genetic structure (PCA) and Redundancy analysis (RDA) based on associations between 6 climate, 3 geographic variables for a) genome-wide nuclear data of all 546 samples analyzed, versus b) genome-wide nuclear and c) chloroplast differentiation across parental-types. In each RDA plot, the direction and length of each arrow corresponds with the strength of prediction (i.e. loading) along each RDA axis. Climate and geographic variables include MAT (mean annual temperature), TD (continentality), PAS (precipitation as snow), RH (relative humidity), MAP (mean annual precipitation), CMD (climate moisture deficit), latitude, longitude, and elevation. Patterns between chloroplast and nuclear DNA are congruent with the exception of one sample (ID= 757_S202_L004) which has *interior P. trichocarpa* nuclear genetic background with *P. balsamifera* chloroplast identity.

Because geographic and climatic predictor variables are often correlated, we performed variance partitioning to tease apart the respective contributions of climate, geography, and their interactions to genomic variance. We observed a large, confounded effect (58.69%) between the climatic and geographic variables chosen for analysis. Removing confounded effects through conditioning revealed climate was a stronger predictor of genetic variance. Climate, conditioned on geography, accounted for 27.28% of explainable genetic variance, whereas geography, conditioned on climate, accounted for just 14.03% of explainable genetic variance (Supplementary.S1; Table S2).

Using the same suite of analyses to investigate associations between environment and ancestry for the 185 parental-types revealed higher contributions of climate and geography to explainable genetic structure. From side-by-side comparisons of nuclear and chloroplast genetic structure in the PCA, we observed similar clustering (Figure 2). Considering the differentiation along PC1 axis, *coastal P. trichocarpa* and *interior P. trichocarpa* appear less diverged based on nuclear data when compared to chloroplast. Similar clustering and environmental associations were also observed in the RDA (Figure 2) and aligned with the associations presented for the 546 samples (Figure 2a). However, with hybrids removed from the analysis, the explanatory variance increased (adj. *r*^2^ = 7.8%; PVE = 51.65%). The explanatory variance based on chloroplast structure was the highest observed (adj. *r*^2^ = 30.1%; PVE = 66.26%). Variance partitioning was also conducted with the hybrids excluded, and with the confounding effects of geography removed, climate explained approximately 40-42% of the genetic variance in both the chloroplast and nuclear data. Conversely, after conditioning on climate, geography explained only 9.46% of the nuclear genetic variance and 2.87% of the chloroplast genetic variance (Supplementary.S1, Table S3).

### Gene flow dynamics shaped by physical barriers

Using departures from isolation-by-distance (IBD) to indicate barriers to gene flow, EEMS analysis identified the continental divide of the Rocky Mountains as a large barrier between *P. trichocarpa* and *P. balsamifera*. Additional barriers to gene flow were identified between transects associated with elevational gradients (Figure 3). For example, *interior P. trichocarpa* within the most southern transect was observed in a region of high local connectivity but was isolated from neighboring individuals identified as *coastal P. trichocarpa* and *P. balsamifera* in other analyses. The EEMS results indicated both Alaska transects were more isolated from the other northern transect, Cassiar, than expected based on IBD. In contrast, individuals spanning *coastal P. trichocarpa* parental-type and hybrids across the Chilcotin, Jasper, and Crowsnest transects were more similar than expected under IBD, suggesting substantial intraspecific migration across latitudes within the southern range of this hybrid zone. This pattern was not observed for *P. balsamifera,* as only individuals located east of the Rocky Mountains exhibited evidence of greater effective migration than expected.

**Figure 3.**
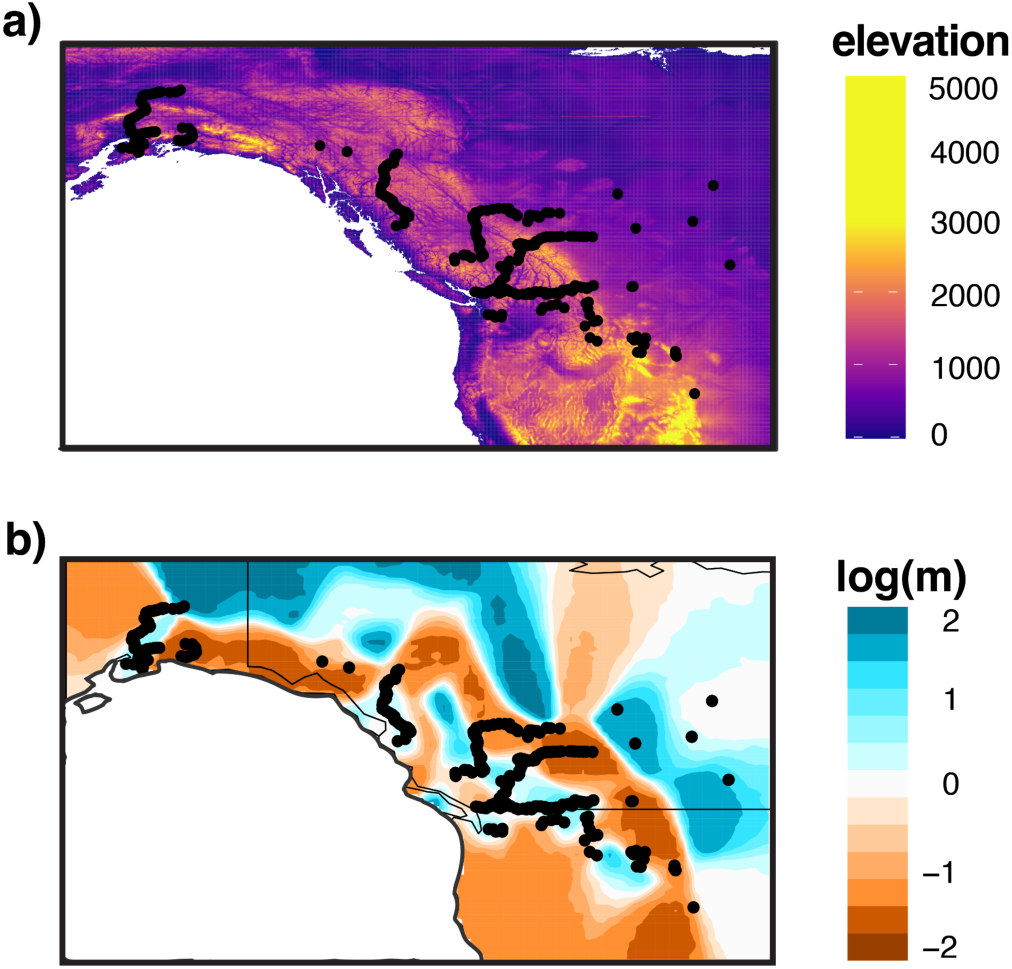
Estimated effective migration surfaces (EEMS) analysis of the relationship between genetic distance and geographical distance for 546 sampled trees across seven transects of the *P. trichocarpa* and *P. balsamifera* hybrid zone to visualize potential barriers to gene flow. Panel a) is a raster map of elevational differences (in meters) across the studied region, and panel b) illustrates putative barriers to gene flow within and among sampled transects. The black dots represent the geographic location of each of the 546 samples. Regions with negative migration rate predictions (brown colors on the log 10 scale) indicate lower genetic connectivity (gene flow) than expected. Regions with positive migration rate predictions (blue colors on the log 10 scale) indicate higher genetic connectivity than expected.

### Demographic inference reveals periods of interspecific vicariance, gene flow, and the emergence of a hybrid lineage

Of the four demographic models of connectivity and divergence evaluated (Table 1b), the model with the greatest support inferred an initial divergence between *coastal P. trichocarpa* - *P. balsamifera* approximately 2.12 million years ago (95% CI: 2.11 – 2.13 million years ago). Following initial divergence, limited gene flow (*m*A ∼ 1.90e-04) continued until approximately 860,000 years ago when gene flow ceased, during which time the two lineages were isolated (Figure 4). Following approximately 96,000 years of isolation, the two lineages experienced secondary contact and 764,000 years ago formed the *interior P. trichocarpa* lineage via admixture. During the initial formation of *interior P. trichocarpa* the proportions of each progenitor lineage were inferred to be approximately 34.8% *coastal P. trichocarpa* (parameter *f*; Table 2) and 65.2% *P. balsamifera* (1 - *f*). Following secondary contact, asymmetrical gene flow was inferred until present, with the highest rates of gene flow observed between *coastal P. trichocarpa* and *interior P. trichocarpa* (*m13* ∼ 5.08e-05 and *m31* ∼ 4.58e-05; Table 2), with a slightly greater rate of genetic exchange from *interior* into *coastal* (*m13*). Gene flow appeared to be strongly asymmetrical, with more movement from *P. balsamifera* into *coastal* and *interior P. trichocarpa* (*m12* ∼ 3.64e-05, *m32* ∼ 3.13e-05, respectively) and less *coastal* or *interior P. trichocarpa* movement into *P. balsamifera* (*m21* ∼ 1.30e-05 and *m23* ∼ 9.08e-06; Table 3). The highest effective population size inferred was for *P. balsamifera* (N_2_ ∼ 100,000) followed by *interior P. trichocarpa* (N_3_ ∼ 53,800) and *coastal P. trichocarpa* (N_1_ ∼ 36,500). The residuals between the model and data fit were small and evenly distributed, suggesting good model fit (Supplementary.S1, Figure S5). In addition, confidence intervals for each parameter were tightly bound to the estimations inferred (Table 2).

**Figure 4.**
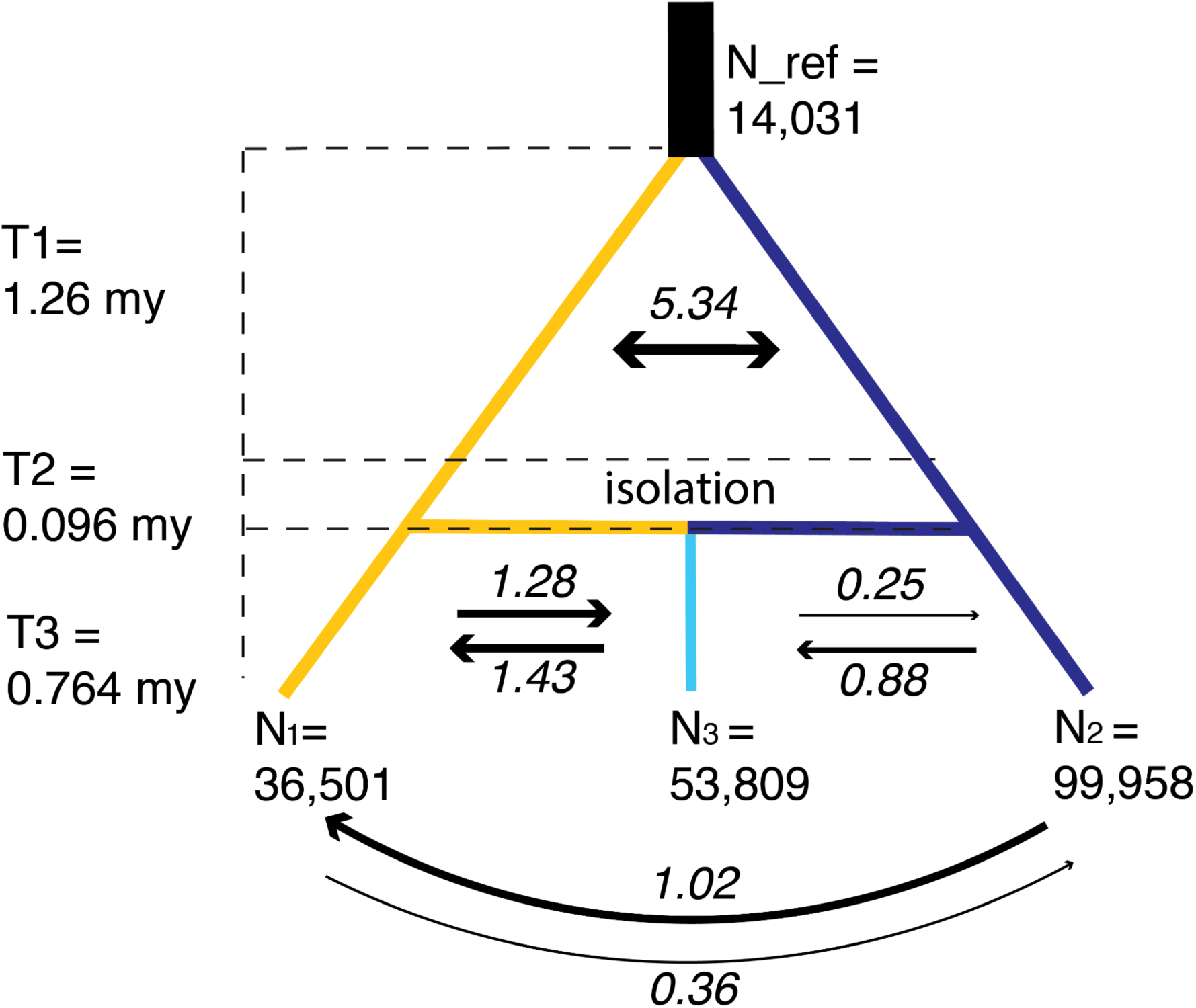
Distribution of samples used in three population demographic inference (a) and the best fit model of divergence among *coasta*l *P. trichocarpa* (in yellow), *P. balsamifera* (in dark blue), and *interior P. trichocarpa* (in light blue). Parameter estimates and confidence intervals for the 3-population model are provided in Table 2. Note that T1, T2, and T3 are time intervals in millions of years (my).

**Table 2.**
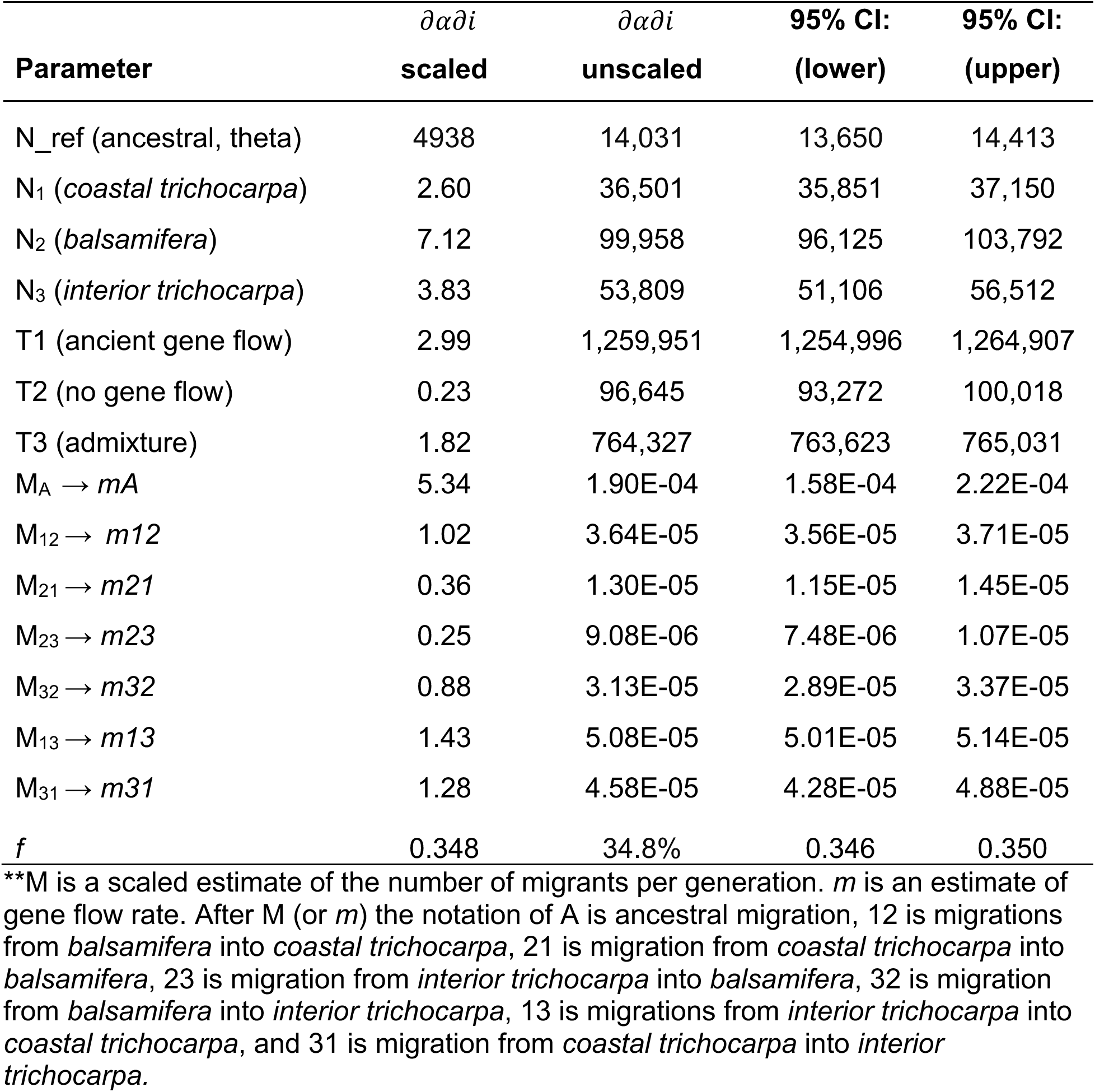
Parameter estimates and 95% confidence intervals (CI) associated with the best fit 3 population demographic model (Topology 3 - T3_IM_SC_IM). Calculations used to unscaling parameter estimates (using theta, mutation rate per generation, and genome length) to meaningful units are provided in Supplementary S3. Each value for T (time) is an interval. Interpretation of each migration (gene flow) parameter is provided in the footnote of this table. The parameter *f* is an estimate of the proportion of *coastal P. trichocarpa* in the formation of *interior P. trichocarpa* at the start of T3. The proportion of *P. balsamifera* is thus 1 - *f*.

| Parameter | $\partial\alpha\partial i$<br>scaled | $\partial\alpha\partial i$<br>unscaled | 95% CI:<br>(lower) | 95% CI:<br>(upper) |
| --- | --- | --- | --- | --- |
| N_ref (ancestral, theta) | 4938 | 14,031 | 13,650 | 14,413 |
| N<sub>1</sub> (*coastal trichocarpa*) | 2.60 | 36,501 | 35,851 | 37,150 |
| N<sub>2</sub> (*balsamifera*) | 7.12 | 99,958 | 96,125 | 103,792 |
| N<sub>3</sub> (*interior trichocarpa*) | 3.83 | 53,809 | 51,106 | 56,512 |
| T1 (ancient gene flow) | 2.99 | 1,259,951 | 1,254,996 | 1,264,907 |
| T2 (no gene flow) | 0.23 | 96,645 | 93,272 | 100,018 |
| T3 (admixture) | 1.82 | 764,327 | 763,623 | 765,031 |
| M<sub>A</sub> → $m_A$ | 5.34 | 1.90E-04 | 1.58E-04 | 2.22E-04 |
| M<sub>12</sub> → $m_{12}$ | 1.02 | 3.64E-05 | 3.56E-05 | 3.71E-05 |
| M<sub>21</sub> → $m_{21}$ | 0.36 | 1.30E-05 | 1.15E-05 | 1.45E-05 |
| M<sub>23</sub> → $m_{23}$ | 0.25 | 9.08E-06 | 7.48E-06 | 1.07E-05 |
| M<sub>32</sub> → $m_{32}$ | 0.88 | 3.13E-05 | 2.89E-05 | 3.37E-05 |
| M<sub>13</sub> → $m_{13}$ | 1.43 | 5.08E-05 | 5.01E-05 | 5.14E-05 |
| M<sub>31</sub> → $m_{31}$ | 1.28 | 4.58E-05 | 4.28E-05 | 4.88E-05 |
| $f$ | 0.348 | 34.8% | 0.346 | 0.350 |

\*\*M is a scaled estimate of the number of migrants per generation. $m$ is an estimate of gene flow rate. After M (or $m$ ) the notation of A is ancestral migration, 12 is migrations from *balsamifera* into *coastal trichocarpa*, 21 is migration from *coastal trichocarpa* into *balsamifera*, 23 is migration from *interior trichocarpa* into *balsamifera*, 32 is migration from *balsamifera* into *interior trichocarpa*, 13 is migrations from *interior trichocarpa* into *coastal trichocarpa*, and 31 is migration from *coastal trichocarpa* into *interior trichocarpa*.

## Discussion

This study provides valuable insights into how complex spatial and temporal dynamics influence hybridization, highlighting the role neutral processes can play in shaping the genetic structure of hybrid zones. A dynamic history between glacial and interglacial cycles suggests that secondary contact over time has had a substantial influence on hybrid zone structure. We identified three distinct genetic clusters geographically distributed across our seven repeated zones of species overlap. Differentiation within and across nuclear and chloroplast genomes supported the presence of three genetic lineages, with tests of topology suggesting one genetic lineage reflects an ancient admixture event between *coastal P. trichocarpa* and *P. balsamifera* leading to the origin of a new hybrid lineage, *interior P. trichocarpa*. While physical geographic barriers have played a significant role influencing historical and contemporary asymmetrical genetic exchange, our results highlight the important role climate gradients alongside geography have played influencing hybrid zone structure. Together, this suggests that genetic patterns observed could be sensitive to climatic shifts, which has implications for understanding the future of these contact zones in a changing climate.

### Demographic shifts associate with shifts in historical climate

According to our best fit 3D demographic model, initial divergence between *P. balsamifera and coastal P. trichocarpa (∼* 2.12 million years ago) aligns with a shift from the Pliocene to the Pleistocene Climate Era (Tan et al., 2018). This initial divergence was inferred to occur amid recurrent gene flow for approximately 1.25 million years, followed by a period of isolation lasting ∼ 96,000 years. Interestingly, this period of isolation not only aligns with the Mid-Pleistocene Transition, when glacial cycles became longer (from ∼40,000 to ∼100,000 years long) and shifts to interglacial climates become abrupt and extreme, but it also coincides in timing with a glacial maximum (∼800,000 years ago; (Chalk et al., 2017; Ellis & Palmer, 2016; Willeit et al., 2019). Secondary contact between *P. balsamifera* and *coastal P. trichocarpa*, along with the origin of *interior P. trichocarpa,* was estimated to have occurred ∼764,000 years ago and coincides with a transition to interglacial climate (Ellis & Palmer, 2016). This suggests opportunities for genetic exchange were regained following expansion from isolated refugia.

While climate has continued to oscillate since the inferred secondary contact event, which might be expected to reduce opportunities for gene flow given the different climatic affinities of each lineage, our demographic model suggested continued genetic exchange. One explanation may be that species distributions have moved in tandem with climate such that populations were proximal enough to maintain gene flow, resulting in a stable hybrid zone. Indeed, the long-distance dispersal of seed and pollen in *Populus* and other forest trees blurs definitions for how geographically isolated species need to be to disrupt gene flow. An alternative explanation may be that periods of strict isolation did occur during extreme climatic shifts but were not sustained long enough for genomic patterns of divergence associated with allopatry to develop, or patterns were broken down following subsequent contact. This latter possibility may characterize hybridizing species with porous genomes whose species boundaries vary temporally with repeated cycles of reinforcement and homogenization (e.g., Mix-Isolation-Mix hypothesis; He et al., 2019) or due to changes in the strength and direction of selection over historically varying climatic optima (e.g., Shang et al., 2020). Among the divergence histories inferred for forest tree species, there is growing evidence for divergence with recurrent gene flow despite abrupt shifts in climate (e.g., Bolte et al., 2022; He et al., 2019; Lexer et al., 2006; Menon et al., 2018).

### Gene flow asymmetries in the context of future climate change

Gene flow was nearly three times higher from *P. balsamifera* into *coastal P. trichocarpa* than from *coastal P. trichocarpa* into *P. balsamifera.* This observation is corroborated by previous estimates of genetic exchange in this hybrid zone by Suarez-Gonzalez et al. (2018), who noted asymmetrical introgression from *P. balsamifera* into *P. trichocarpa*. This asymmetry may suggest *P. balsamifera* has a demographic advantage within the ecotonal transition (e.g., dispersal down an elevational gradient or with river flow from inland to coast). On the other hand, lower rates of introgression between *P. balsamifera* and *interior P. trichocarpa* may be indicative of stronger geographical isolation, a lack of hybrids geographically positioned between these genetic clusters to facilitate backcrossing, or even more likely, asynchronous reproductive phenology. According to descriptions within (Haeussler et al., 1990) flowering times for interior populations of *P. trichocarpa* can be nearly one month later than those reported for populations of *P. trichocarpa* (coastal) and *P. balsamifera* in British Columbia, Canada.

Gene flow between *coastal P. trichocarpa* and *interior P. trichocarpa* were greatest relative to all inferred between-lineage contemporary estimates (Figure 4b). Indeed, *interior P. trichocarpa* is genetically more similar to *coastal P. trichocarpa* based on patterns of nuclear differentiation. However, despite this, the demographic model that received the greatest support suggested that in the formation of *interior P. trichocarpa* (∼764,000 years ago) only 35% of the hybrid genome reflected ancestry from *coastal P. trichocarpa*. Taken together, these results may indicate that higher rates of contemporary gene flow between *coastal* and *interior P. trichocarpa* homogenized population differences following the initial divergence of the *interior P. trichocarpa* lineage. Regardless, the influx of new genetic recombinants through introgression is evident across all three parental-types and may impact adaptation to a changing climate (Aitken et al., 2008; N. Barton & Bengtsson, 1986; Martinsen et al., 2001).

### Climate and landscape heterogeneity contribute to genetic structure

Landscape-induced restrictions to gene flow have contributed to patterns of differentiation across the hybrid zone. Estimated effective migration surfaces (EEMS) suggest gene flow has likely been suppressed by elevational barriers through the Rocky Mountains of northwestern North America. West of the Rocky Mountains, we observed variation in effective migration rates within the *P. trichocarpa* distribution described by (EL Little Jr., 1971). For example, effective migration was greater within the northern transects of Alaska and Cassiar but not between these two transects. Similarly, there appeared to be substantial gene flow between hybrids within the ‘mid-latitudinal’ transects (Chilcotin, Jasper, and Crowsnest), but migration was restricted along the coastal regions of these three transects where parental-types of *coastal P. trichocarpa* were identified. This suggests gene flow between *P. trichocarpa* populations is lower than expected based on expectations of isolation-by-distance. Thus, landscape barriers to gene flow likely contribute to strong population structure observed within the range of *P. trichocarpa* (Geraldes et al., 2014; Slavov et al., 2012).

Redundancy analysis (RDA) and variance partitioning revealed genetic structure and climate are significantly associated across the hybrid zone. The Rocky Mountains present environmental transitions from drier and colder conditions on the eastern side (favoring *P. balsamifera*) to more maritime climates on the western side of the Rockies (favoring *coastal P. trichocarpa*). The distributions and climate associations we observed between *P. balsamifera, coastal P. trichocarpa*, and *interior P. trichocarpa* (Figure 2a) are comparable to other hybrid zones within the region, including *Picea glauca, P. sitchensis*, and *P. engelmannii* hybrid zones (De La Torre et al., 2015; Hamilton et al., 2013b, 2015). In both systems, maritime to continental climate transitions were associated with genomic ancestry. Because the frequency of hybrid genotypes is largely associated with ecotonal transitions, this suggests these hybrid zones likely reflect a model of bounded hybrid superiority. However, quantifying fitness in reciprocal transplant experiments is needed to test the bounded hybrid superiority model.

The admixed lineage, *interior P. trichocarpa*, appears to be bounded to a drier and warmer climate than either of its progenitors. The genetic structure of *interior P. trichocarpa* is associated with high climate moisture deficit. If selection maintains genetic structure among the lineages observed with our study, we may find trait divergence, especially those underlying water-use, across genetic lineages. Interestingly, *Populus* hybrid lineages of the Qinghai-Tibet Plateau (QTP) also inhabit landscapes with higher aridity relative to their progenitors. However, survival and growth measurements within common garden experiments suggested phenotypic plasticity may have contributed to adaptive divergence and persistence in environments beyond the parental species’ range (Shi et al., 2024). A similar experimental design could be employed for *coastal P. trichocarpa, interior P. trichocarpa,* and *P. balsamifera* to test if the admixed lineage (*interior P. trichocarpa*) exhibit transgressive traits relative to parents while also evaluating the role of plasticity to success in novel environments. In the absence of trait data, we can leverage genomic data to differentiate between, and functionally assess, genomic regions associated with restricted or excess introgression that may underlay adaptive divergence.

### Nuclear and chloroplast genomes provide complementary insights into hybrid zone dynamics

The estimates observed herein suggest 2.11 - 2.13 million years ago *P. trichocarpa* and *P. balsamifera* diverged and are congruent with observations for this species-pair from (Sanderson et al., 2023). The differentiation we observed in the chloroplast not only aligns with patterns of nuclear differentiation among the three assigned parental-types but also supports the observations and interpretations of previous chloroplast assessments of divergence in *Populus*. (D. I. Huang et al., 2014) ascribed the deep chloroplast divergence of *P. trichocarpa* from *P. balsamifera* to chloroplast capture of an ancient, now extinct, lineage. While the genome-wide patterns of nuclear differentiation (e.g., PCA, *F*_ST_) suggests *coastal P. trichocarpa* and *interior P. trichocarpa* are more closely related, our phylogenetic inference from the chloroplast placed *coastal P. trichocarpa* outside the clade of *interior P. trichocarpa* and *P. balsamifera*. Only one individual out of 185 did not have a matching chloroplast - nuclear parental-type assignment. Considering our demographic modeling and the fact that our focal taxa exhibit dioecy with maternal inheritance of the chloroplast genome, the origin of *interior P. trichocarpa* likely resulted from admixture between *P. balsamifera* seed parents and *coastal P. trichocarpa* pollen parents. Given the inferred length of time that has passed since the admixed origin (∼764,000 years), the *interior P. trichocarpa* chloroplast genome has likely evolved following divergence. Co-evolution and co-adaptation between the chloroplast and nuclear genomes may have contributed to chloroplast divergence of *interior P. trichocarpa* from the original seed parent (Greiner & Bock, 2013) and would be an interesting future avenue for research.

## Conclusions

Here, we provide insight into the evolutionary processes shaping divergence and hybridization across space and time in a *Populus* system of western North America. Demographic models support the origin of a hybrid lineage following secondary contact of ancestral lineages and a history of asymmetrical gene flow between focal taxa. Our results highlight the influence of climate and geography to genetic structure and divergence, as well as the influence of landscape-level attributes to gene flow dynamics. Overall, our study sheds light on the role demographic processes can play within a long-lived hybrid zone across space and time and highlight the need for fine-scale assessment of regions of the genome important to introgression and adaptation for species management in a changing climate.

## Acknowledgements

We thank Joe Braasch, Lionel Di Santo, Sonia DeYoung, Sara Klopf, Jack Woods, Ben Woods, Raju Soolanayakanahally for assisting with field sample collection. We also thank Kyle Peer, Clay Sawyers, and Deborah Bird from the Virginia Tech Reynold’s Homestead Forestry Research Station for their assistance with plant propagation. We thank Ryan Gutenkunst and Andrew Eckert for advice and communications related to demographic inference. We acknowledge the valuable questions from and conversations with our team members during project calls and meetings: Cigdem Kansu, Susanne Lachmuth, Baxter Worthing, and Alayna Mead.

## Author contributions

MCF, JH, SRK, and JAH conceived the study and collected the samples. MC extracted DNA. TP processed genomic data. CEB analyzed genomic data, conducted demographic inference, and performed genetic-environment associations. MZ and BS assembled chloroplast genomes and performed chloroplast analyses. SRK, JH and JAH advised analyses. CEB and JAH led the writing of the manuscript. SRK, JH, and MCF provided valuable feedback.

## Data availability

Sequence reads for the 546 samples analyzed in this manuscript have been archived on NCBI (PRJNA996882). Chloroplast genomes are archived under BankIt2793276 (Accession numbers PP274128 - PP274690). Both nuclear and chloroplast sequences will remain under embargo until publication. Scripts for variance partitioning, RDA, and demographic inference can be found at https://github.com/boltece/Poplar_divergence.

## Supporting Information

**Supplementary.S1 -** Supplementary methods (appendices) and figures

**Supplementary.S2 -** Sample summaries for climate and geographic data, admixture coefficients, and PCA and RDA coordinates

**Supplementary.S3 -** Summarized results from demographic inference including AIC calculations, unscaling of *∂α∂i* parameter estimates, and confidence interval calculations

