## Supplementary.S1 for "Genomic insights into hybrid zone formation: the role of climate, landscape, and demography in the emergence of a novel hybrid lineage between *Populus trichocarpa* and *P. balsamifera*"

**Appendix 1. Protocol for propagation of vegetative branches under greenhouse conditions**

Vegetative branch cuttings from 576 trees were provided a unique identifier and latitude, longitude, and elevation of origin were recorded (Supplementary.S2). Vegetative branch cuttings were divided and propagated at Virginia Tech. Each cutting was exposed to a 30 second Zerotol 2-0 (27.1% hydrogen peroxide, 2% peroxyacetic acid) fungal dip treatment at 1:100 ratio of Zerotol 2-0 to water. Cuttings were then dipped in Garden Safe Take Root hormone, rooted into a standard rooting mix (peat moss, vermiculite, perlite, osmocote, bone meal, gypsum, epsom salt, and dolomite), and subsequently placed on a mist bench (Phytotronics Nova mist controller, DGT blue/white mist .53 GPM). Mist was applied for 38 days (March 9 - April 17, 2020); for the first 31 days, every 10 minutes for 8 seconds, then the last 7 days, every 60 minutes for 10 seconds. Greenhouse conditions included a day temperature of 24 °C, a night temperature of 16 °C without supplemental lighting. For each of the 576 samples, approximately 100mg of young leaf tissue was weighed and frozen using liquid nitrogen, ground into fine powder with mortar and pestle, and transferred into a microcentrifuge tube for genomic DNA extraction.

**Appendix 2. Extractions, library preparation, and variant calling protocols for whole genome resequencing data**

### DNA was extracted from all samples using the Qiagen plant DNeasy kit, with one modification. In place of a QIAshredder column, a phenol-chloroform extraction was used. For samples with very low DNA concentration, a second extraction was performed using cetyltrimethylammonium bromide (CTAB). DNA concentrations and quality were quantified using the Qubit dsDNA HS Assay kit and Nanodrop spectrophotometer. Genomic DNA libraries were constructed at the Duke University Center for Genomic and Computational Biology, using an Illumina DNA Prep kit (Illumina Inc., San Diego, USA). Genomic libraries were sequenced on an S4 flow cell in 2x150bp format on an IlluminaNovaSeq 6000 instrument with 64 samples per lane. De-indexing, QC, trimming adapter sequences, and sequence preprocessing were completed by the sequencing facility.

### Subsequent bioinformatic analysis was performed on Virginia Tech’s Advanced Research Computing System (ARC) using Burrows-Wheeler Aligner (BWA) to map reads to the *P. trichocarpa* reference genome (v4.0), and the resulting SAM files were converted to BAM format with SAMtools (Li and Durbin 2010). The Genome Analysis Toolkit (v3.7) Haplotype Caller algorithm was then used to generate individual gVCF files, which were merged in a single VCF file with the GATKGenotype GVCFs function. This raw VCF had ~82 million variant calls and was quality-filtered for variants that had poor map quality (MQ < 40.00), elevated strand bias (FS > 40.000, SOR > 3.0), differential map quality between reference and alternative alleles (MQRankSum < -12.500), positional bias between reference and alternate alleles (ReadPosRankSum < -8.000), or low coverage depth (QD < 2.0).

Variant calling for the chloroplast (cp) data was performed by using paired-end Illumina reads of the 185 individuals (i.e., parental-type samples) using bcftools 1.14 (Li 2011). The *P. trichocarpa* cp genome (NC_009143.1) was chosen as a reference, but prior to mapping, one of the inverted repeat regions was manually removed because the duplicated regions in cp genomes can hamper alignments. Reads were aligned to the modified *P. trichocarpa* cp genome using Bowtie and stored as SAM files using Samtools 1.15 (Li et al. 2009). The individual SAM files were then converted to BAM files and sorted for efficient processing. Subsequently, variant calling was performed on the individual sorted BAM files and merged using bcftools. The VCF file was filtered with a minimum variant quality threshold (QUAL) of 30, maximum missing data of 0.05, and indels were removed. A total of 2,848 variants were kept after filtering. A PCA was conducted using the same approach described in the main text for nuclear data (Patterson et al. 2006).

**Appendix 3. Ancestry assignment within a *Populus* species complex**

### In select southern parts of our sampling region, *Populus trichocarpa* and *P. balsamifera* co-occur with *Populus angustifolia*. To account for potential sampling of non-focal poplar species and remove samples with *P. angustifolia* ancestry, we merged the genetic data for our 575 samples with genetic data available for 48 samples of *P. angustifolia* (Chhatre et al. 2018; NCBI SRA study- SRP070954). Across the 48 samples of *P. angustifolia*, we identified 94,278 biallelic SNPs with minor allele count greater than 1 and no more than 30% missing data. After imputing the missing data in beagle for the *P. angustifolia* genetic data, we merged the variant call files in bcftools (version 1.18) which yielded 18,025 SNPs across 19 chromosomes (scaffolds were removed) with minor allele counts greater than 1 and no missing data.

### Genetic structure for 623 samples and 18,025 SNPs was analyzed using the program Admixture (Alexander et al. 2009). No representation of *P. angustifolia* was noted in our field collections of 575 individuals above K=4. However, another ancestral group was collected and hybrids with this unknown group were noted in 29 of the 575 samples (Supplementary.S1, Figure S1). Principal component analysis confirmed the presence of an additional ancestral group (Supplementary.S1, Figure S2). Cluster assignments and PC loadings are provided in Supplementary.S2. For the purposes of this study, we excluded the 29 samples with unknown genomic ancestry, leaving 546 individuals for subsequent genetic analyses.

**Appendix 4. Building the consensus fasta file for ancestral state**

We downloaded whole-genome sequence data for *Populus alba*, *P. davidiana*, *P. tremuloides*, *P. tremula*, and *P. deltoides* (Source data and accession numbers for each sample are summarized in Supplementary_S1, Table S1). Genomes were then each aligned to the *P. trichocarpa* reference using bwa, genotyped using GATK HaplotypeCaller and GenotypeGVCFs, and merged with representative samples of *P. trichocarpa* and *P. balsamifera* from our dataset. After setting sites with depth less than 3 or quality less than 20 to missing, we generated a fasta file for each sample using *bcftools* ‘consensus’, and subsequently generated a multi-species fasta for each chromosome and scaffold of the poplar genome. Using PHAST ‘phylofit’ (Siepel and Haussler 2004) we fit a phylogenetic model to each chromosome and estimated the marginal probability for each base across the genome using PHAST ‘prequel’. The base with the highest marginal probability for each position was then used to encode a fasta representing the state of the common ancestor of *P. balsamifera* and *P. trichocarpa* using PHAST2fasta (Luqman et al. 2023). Polarization of the vcf file was performed with *vcfdo*, version 1.9 (https://github.com/IDEELResearch/vcfdo).

**Table S1.** Source data for consensus-based ancestral state inference

| **ID** | **Species** | **Accession** | **Source** | **Link** | **Notes** |
| --- | --- | --- | --- | --- | --- |
| pal12 | P. alba | CRR330694 | GSA-China | https://ngdc.cncb.ac.cn/gsa/browse/CRA005197 | NA |
| pda25 | P. davidiana | CRR330706 | GSA-China | https://ngdc.cncb.ac.cn/gsa/browse/CRA005197 | NA |
| Alb6-3 | P. tremuloides | SRR2749823 | NCBI | https://www.ncbi.nlm.nih.gov/sra/?term=SRR2749823 | NA |
| asp201 | P. tremula | ERR4842717 | NCBI | https://www.ncbi.nlm.nih.gov/sra/ERX4712504[accn] | NA |
| LAR-12_S22_L001 | P. balsamifera | NA | NA | NA | our data |
| 504_S46_L001 | P. trichocarpa | NA | NA | NA | our data |
| S3239 | P. deltoides | SRR11622684 | NCBI | https://www.ncbi.nlm.nih.gov/sra/SRX8186919[accn] | NA |

**Table S2.** Partitioning of Variance of the full and partial contributions of climatic and geographic variation on 1) nuclear genetic variation across 546 samples of the hybrids zone), 2) nuclear genetic variation across parental-types used in demographic inference, and 3) plastid genetic variation across parental-types used in demographic inference. Respectively, the six climatic predictors and three geographic predictors together explained 38.26%, 51.65%, and 66.26% of the total genetic variation. Adjusted *r*^2^ represents the percent of genetic variance that could be explained by the predictors after removing correlative effects. The proportion of variance explained (PVE) represents the overall contribution of the predictors, both full and partial, to genetic structure. Models are ordered by the greatest PVE.

|  | **genome-wide, nuclear** | | **parental-types, nuclear** | | **parental-types, cpDNA** | |
| --- | --- | --- | --- | --- | --- | --- |
| **Models** | **Adj. r^2^**  **(%)** | **PVE (%)** | **Adj. r^2^**  **(%)** | **PVE (%)** | **Adj. r^2^ (%)** | **PVE**  **(%)** |
| Genetics  ~ Climate + Geo | 2.17 | 38.26 | 7.78 | 51.65 | 30.09 | 66.26 |
| Genetics  ~ Climate \| Geo | 0.59 | 27.28 | 3.24 | 41.63 | 12.24 | 40.68 |
| Genetics  ~ Geo \| Climate | 0.31 | 14.03 | 0.74 | 9.46 | 0.80 | 2.87 |
| Residuals | 97.83 | - | 92.22 | - | 69.91 | - |

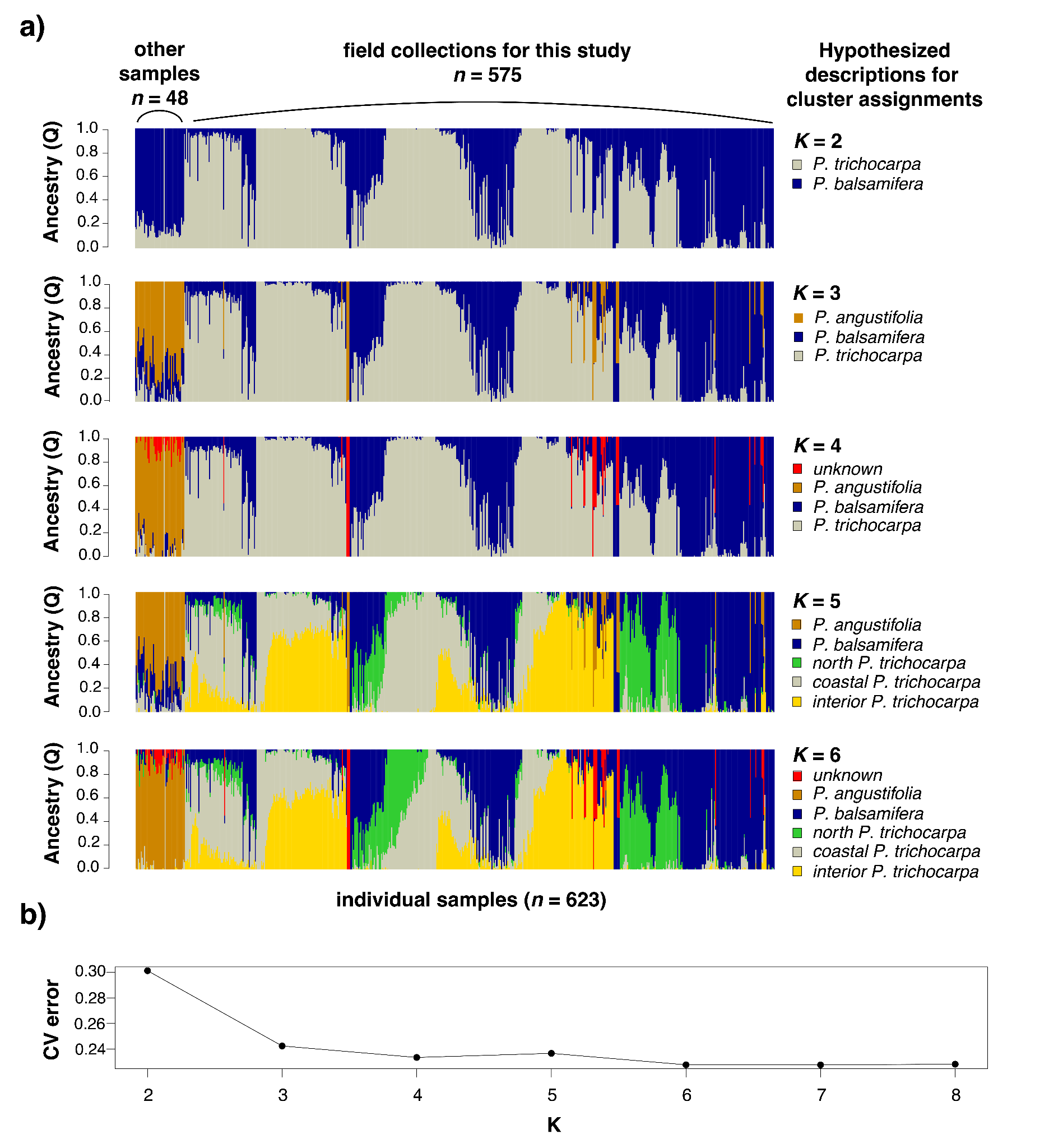

**Figure S1.** Analysis of admixture to determine if other poplar species, aside from the focal species *Populus trichocarpa* and *Populus balsamifera*, were sampled during field collections and subsequently whole genome sequenced. Genetic data included in this analyses (18,025 biallelic single nucleotide polymorphisms) was the result of merging the 575 successfully genotyped samples from whole genome resequencing with 48 samples of *Populus angustifolia* that were previously analyzed in Chhatre *et al.* (2018; NCBI SRA project SRP070954). *Populus angustifolia* is also native to the pacific Northwest. Panel a) illustrates the clusters (*K*) identified in analyses of *K* = 2 through *K* = 6. Each cluster was hypothetically assigned to a species or genetic grouping that aligned with known geographical distributions (Little, 1971). Panel b) provides cross validation errors associated with admixture analysis of *K* = 2 through *K* = 8. The cluster assignment with the lowest cross validation error was *K* = 6, suggesting this assignment best fit the data. In both K = 4 and K = 6, there is a genetic cluster that likely belongs to a different species of poplar. We deemed it “unknown.”

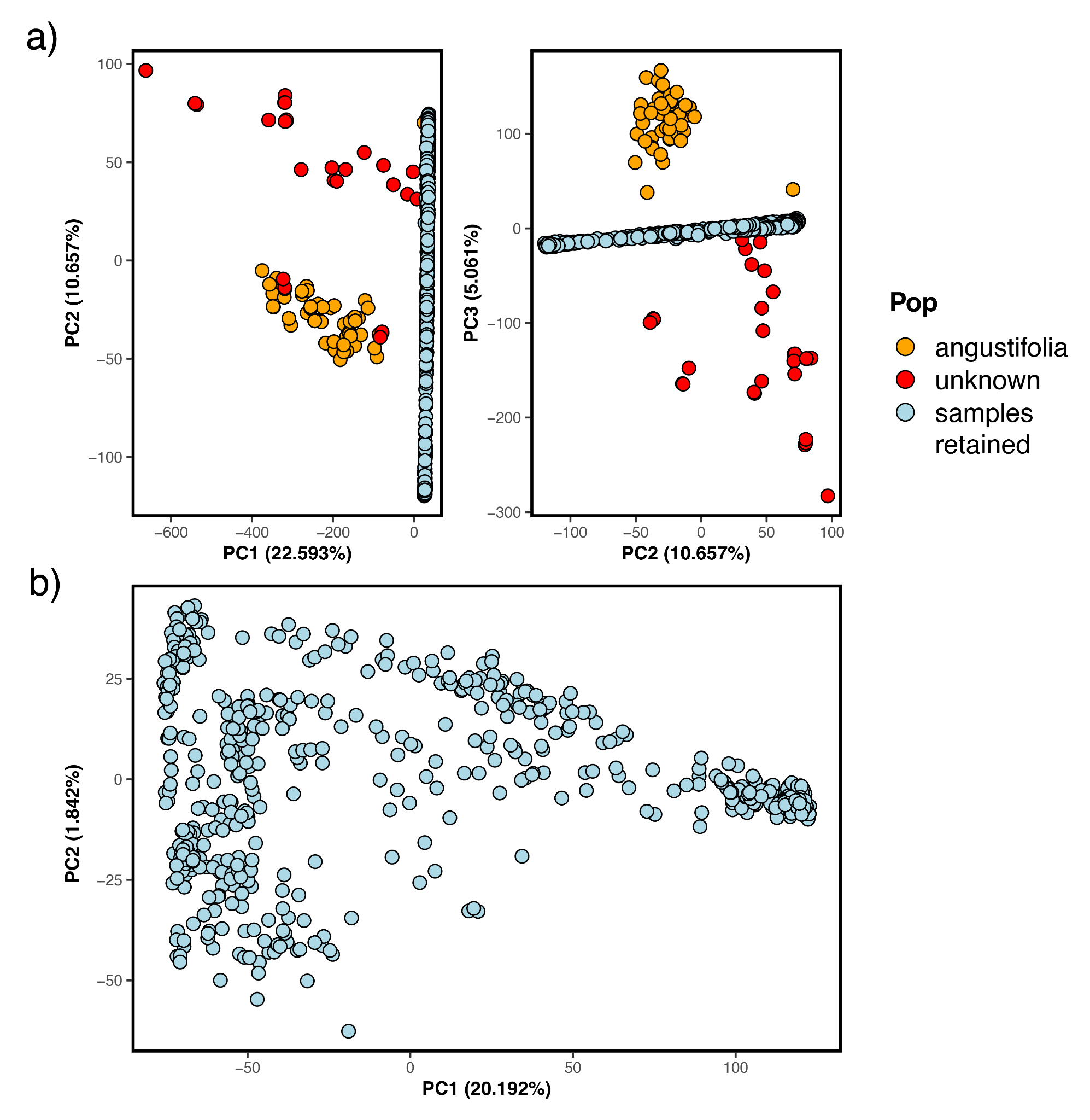

**Figure S2**. Principal Component Analysis based on differentiation across 17,953 SNPs for a) all 575 sampled trees in addition to the 48 *P. angustifolia* samples sourced from Chhatre *et al.* (2018). The unknown (red dots) were first identified as ‘unknown’ through admixture analysis (Figure S1).

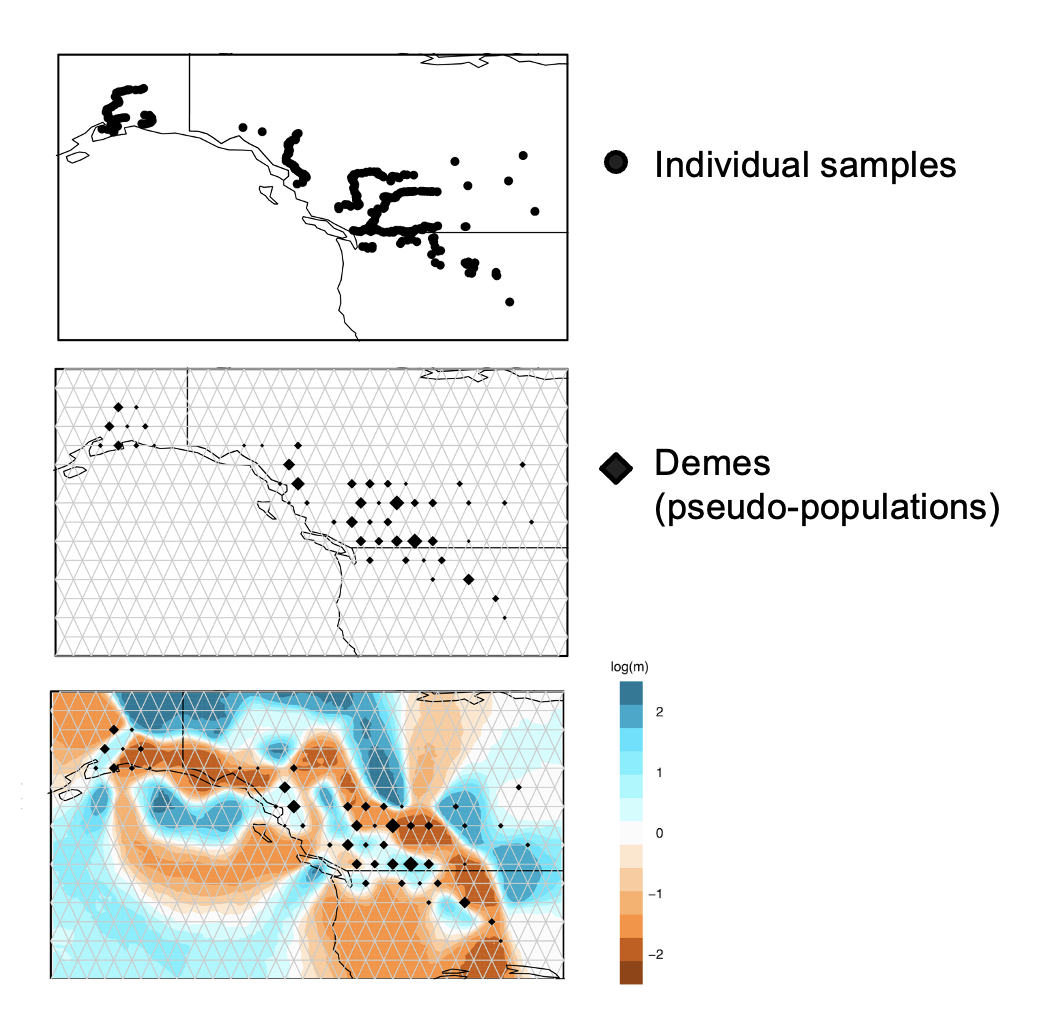

**Figure S3.** Demes (diamonds; second panel) formed from 546 individual samples (black dots; top panel) in a 500 deme grid space (gray lines; second panel) to calculate Estimated Effective Migration Surfaces (connectivity in log(m); bottom panel). Demes were formed based on geographic proximity to a grid vertice. The larger the diamond, the larger the number of individual samples within the deme.

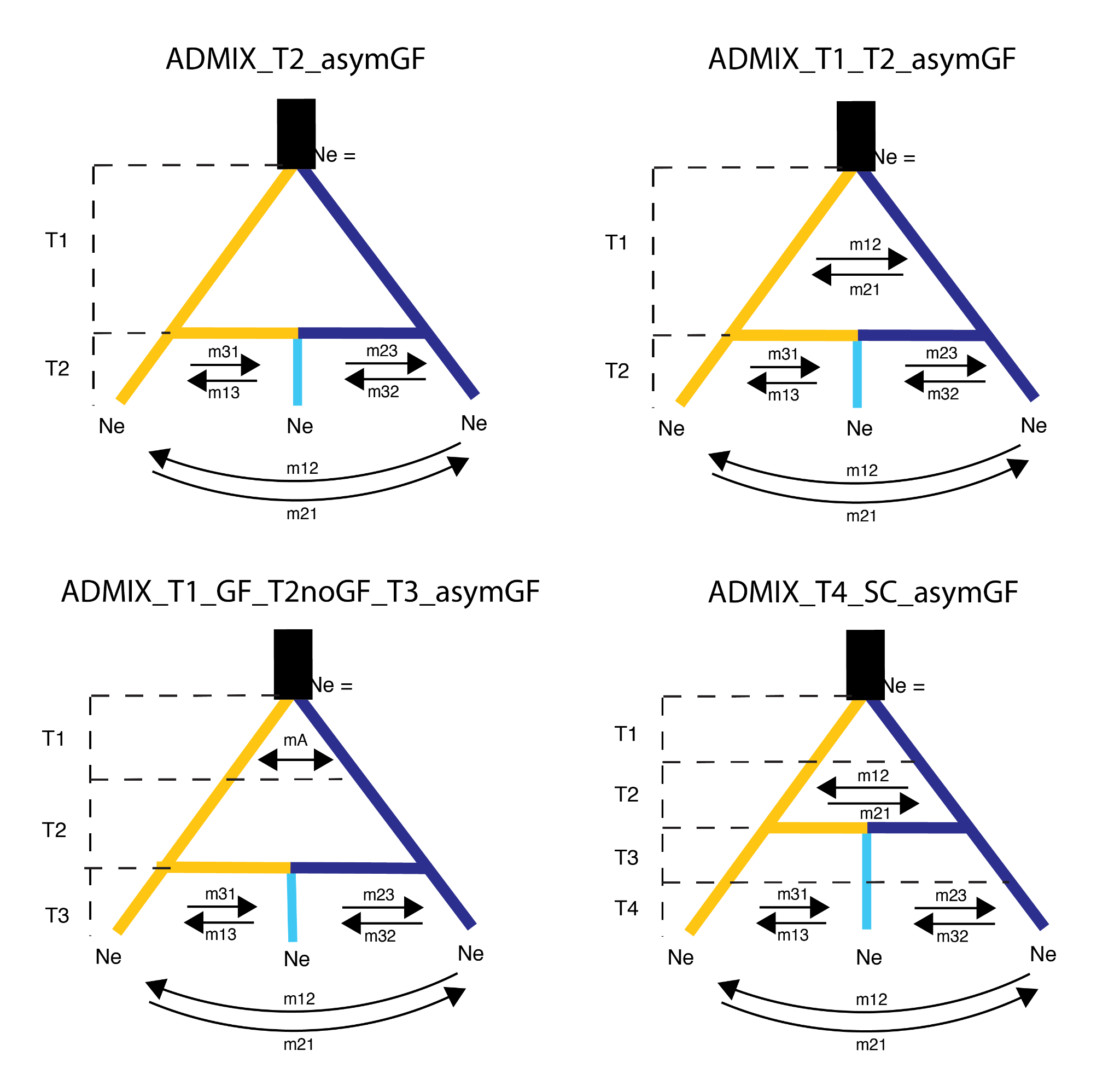

**Figure S4.** Gene flow scenarios during divergence that were tested in $\partial\alpha\partial i$ (v.2.1.1; Gutenkunst *et al.,* 2009).

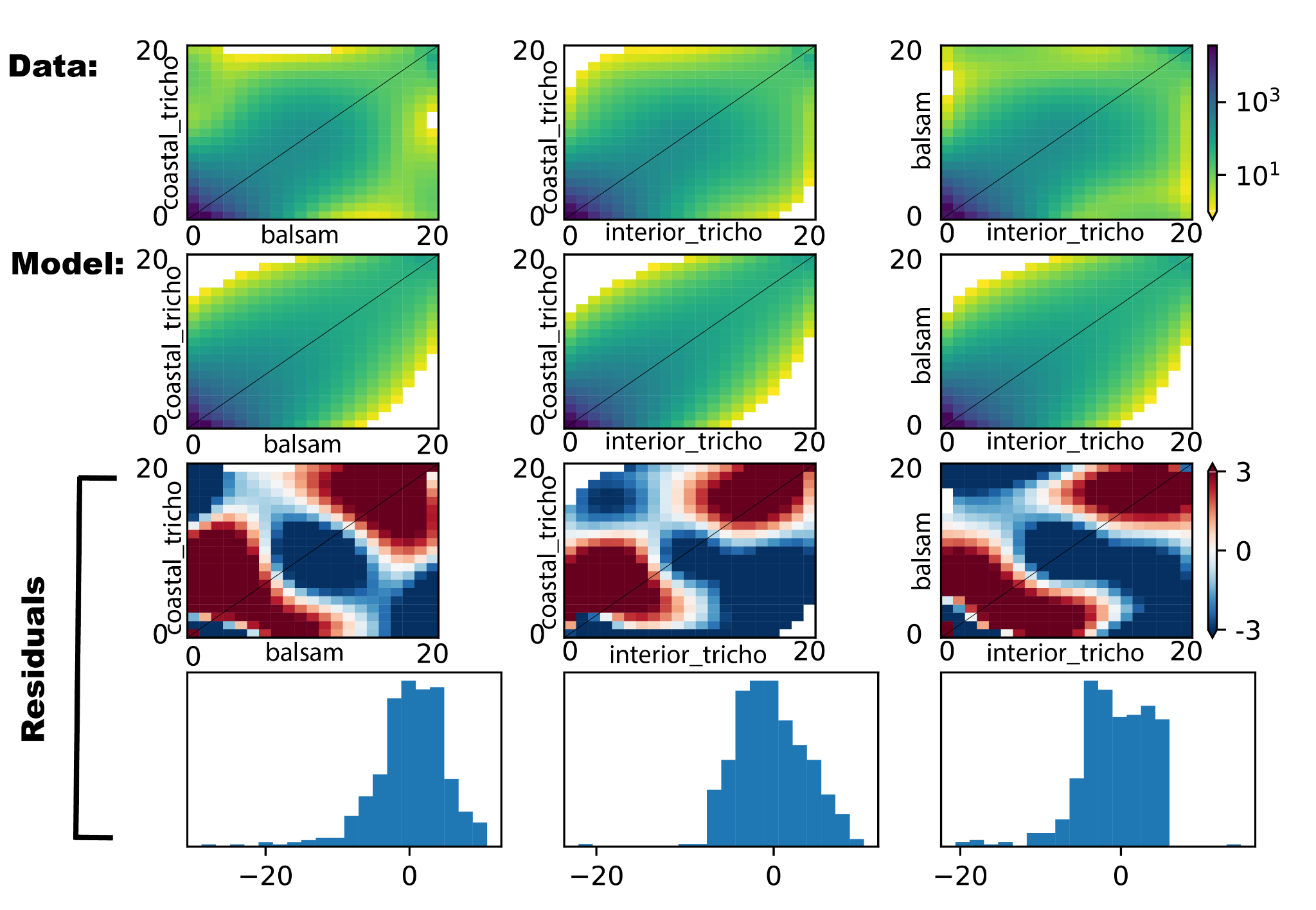

**Figure S5.** Resulting plots of joint site frequency spectrums (jSFS) from downsampling data to 20 chromosomes per population. Top plots of each panel show data to model comparisons of jSFS patterns. Bottom plots of each panel provide quantification and visualization of residuals from the data to model fit.
